# Increased task-dependent functional connectivity by stimulation of the hippocampal network predicts memory enhancement

**DOI:** 10.1101/676643

**Authors:** Kristen N. Warren, Molly S. Hermiller, Aneesha S. Nilakantan, Joel L. Voss

## Abstract

Successful episodic memory involves dynamic increases in the coordination of activity across distributed hippocampal networks, including the posterior-medial network (PMN) and the anterior-temporal network (ATN). We tested whether this up-regulation of functional connectivity during memory processing can be enhanced within hippocampal networks by noninvasive stimulation, and whether such task-dependent connectivity enhancement predicts episodic memory improvement. Participants received stimulation targeting either the PMN or an out-of-network control location. We compared the effects of stimulation on fMRI connectivity measured during an autobiographical memory retrieval task versus during rest within the PMN and the ATN. PMN-targeted stimulation significantly increased connectivity during memory retrieval versus rest within the PMN. This effect was not observed in the ATN, or in either network due to control out-of-network stimulation. Task-dependent increases in connectivity due to PMN-targeted stimulation within the medial temporal lobe predicted improved performance of a separate episodic memory test. It is therefore possible to enhance the task-dependent regulation of hippocampal network connectivity that supports memory processing using noninvasive stimulation.

## Introduction

There is substantial recent interest in treating memory disorders via brain stimulation^1-3^. Episodic memory depends on the hippocampus^4,5^ as well as on the distributed set of regions that form a hippocampal-cortical network^6-10^ comprising distinct anterior-temporal and posteriormedial components^10,11^. The goal of this study was to determine whether noninvasive stimulation targeting the hippocampal-cortical network can enhance network connectivity measured during memory processing, and whether such enhancement is related to episodic memory improvement. We targeted specific portions of the posterior-medial network (PMN) and therefore further hypothesized that stimulation would disproportionately impact the PMN rather than the anterior-temporal network (ATN). This question is of substantial mechanistic and practical significance, given that increased connectivity due to stimulation should manifest primarily during memory processing that depends on the network, and such task-dependent modulation would be essential for any effective intervention intended to improve memory ability.

Invasive stimulation of the hippocampus or its direct mesial-temporal afferents has primarily been associated with memory disruption^12-14^. However, the hippocampal network can serve as a target for memory improvement^15-18^. Noninvasive transcranial electromagnetic stimulation (TMS) targeting cortical network locations has been shown to improve memory^19-24^ and alter hippocampal-cortical network fMRI activity^19-23^, especially within the PMN^21-23^, for durations that substantially outlast the stimulation period. Because stimulation increases connectivity among hippocampal-cortical network regions^22^ and relatively low network connectivity is associated with poor episodic memory^25-30^, it is tempting to hypothesize that memory improvements due to stimulation occur via overall increased connectivity of the hippocampal-cortical network. However, successful memory typically relies on dynamic reconfiguration of fMRI connectivity within the PMN and ATN in response to memory processing demands^31-36^. Thus, stimulation might need to produce task-dependent and location-specific, rather than nonspecific, increases in hippocampal-cortical network connectivity in order to benefit memory.

Indeed, several lines of evidence suggest that memory enhancement should require that hippocampal-cortical network fMRI connectivity increases occur in response to memory processing demands. For instance, effective memory encoding and retrieval is associated with location-specific and task-dependent increases in stimulus-evoked fMRI activity within the network^37-41^. Correspondingly, better memory can be predicted by fMRI connectivity related to specific memory processes, such as recollection^33,35^. In rodents, theta-gamma synchrony in the hippocampus is observed during memory processing, indicating connectivity changes modulated by memory demands^18^. Pharmacological disruption of hippocampal theta-gamma synchrony and memory can be rescued by theta-burst stimulation of the fornix, indicating that this marker of connectivity is critical for memory^18^. Furthermore, amnestic states, such as those caused by hippocampal lesions, are associated not only with reduced hippocampal-cortical network connectivity, but also with increased connectivity between the hippocampal network and other networks^29,30^, suggesting that nonspecific increases in connectivity can be problematic. Demonstration that stimulation can affect memory task-dependent changes in connectivity within specific targeted portions of the hippocampal-cortical network therefore is crucial for its evaluation as an effective memory intervention. This question has not been addressed, as the current standard for assessing the effects of brain stimulation on networks is via fMRI connectivity measured during the resting state^22,42-44^, which by definition is insensitive to task-dependent connectivity.

We evaluated whether stimulation can alter task-dependent fMRI connectivity within the PMN and ATN. One group of participants (n=16) received multi-session, high-frequency TMS targeting the PMN via parietal cortex, which is a robust component of the PMN^10,11^. A separate control group (n=16) received the same TMS regimen targeting a control prefrontal cortex (PFC) location that is not robustly part of the hippocampal-cortical network. Each group also received site-specific sham-control stimulation administered in counterbalanced order with real stimulation (Figure 1). We compared the effects of stimulation on fMRI connectivity within the PMN, ATN, and whole-brain (via an exploratory analysis) measured 24 hours later during a memory retrieval task versus during the resting state. The memory retrieval task involved extended periods of autobiographical memory retrieval, which has been shown to cause robust hippocampal-cortical network connectivity changes primarily in PMN regions compared to the resting state^31^. Thus, we were able to test for task-dependent and network-specific effects of stimulation on fMRI connectivity.

**Figure 1:**
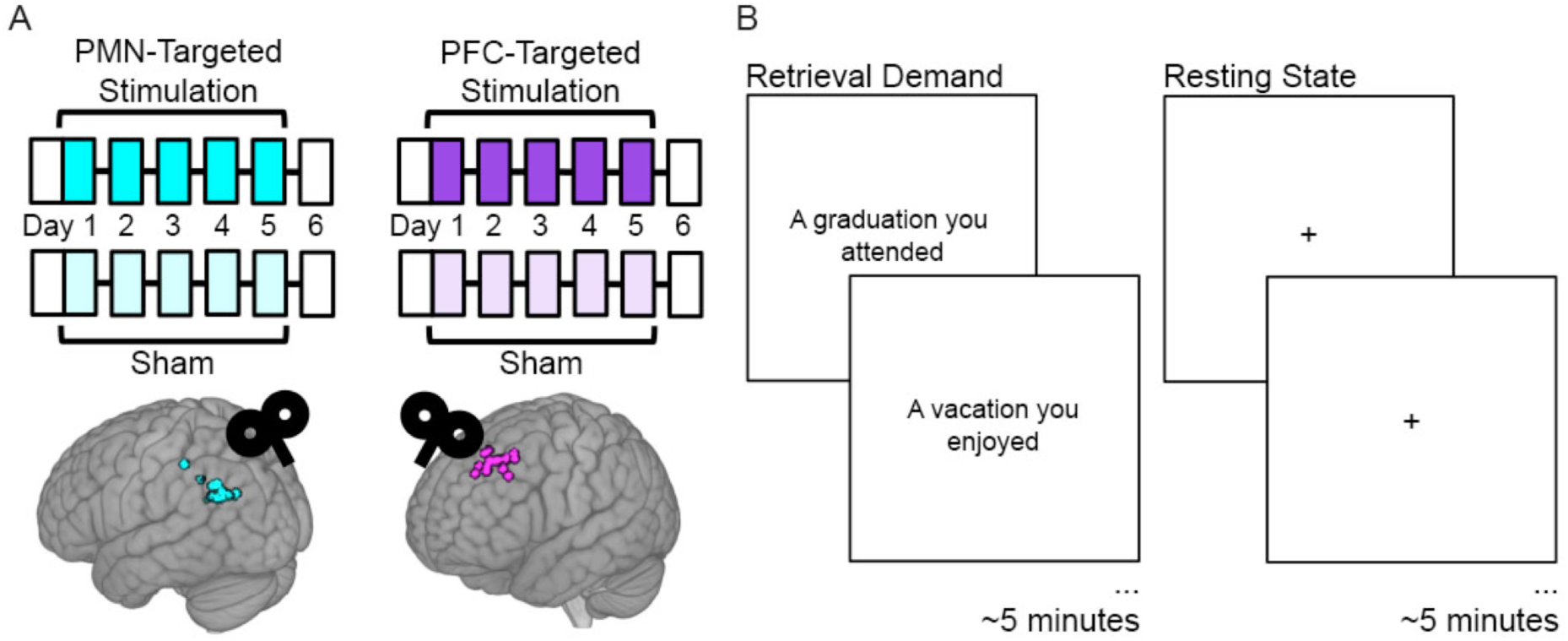
Experiment Design. (A) Subjects received five consecutive daily sessions of high-frequency (20Hz) repetitive TMS delivered to a subject-specific parietal cortex location of the PMN selected based on high resting-state fMRI connectivity with the hippocampus (PMN-Targeted Stimulation). Subjects received real stimulation and sham stimulation during different weeks, in counterbalanced order. Before and ∼24 hours after stimulation, subjects completed fMRI and memory assessments (white boxes). The same procedures were performed for a distinct control group of subjects, but with stimulation delivered to subject-specific locations of out-of-network prefrontal cortex (PFC-Targeted Stimulation). Circles indicate stimulation locations for each participant. (B) fMRI connectivity was measured during the resting state and during an autobiographical memory retrieval task, for which subjects were shown prompts describing common life events and asked to vividly recall personal events matching the prompts.

We hypothesized that PMN-targeted stimulation would increase fMRI connectivity during memory retrieval as compared to the resting state relative to sham-control stimulation, whereas out-of-network PFC-targeted stimulation would not. We additionally hypothesized that these connectivity changes would be specific to the PMN, as it was targeted, contributes to autobiographical memory retrieval^10^, and responds to the stimulation regimen that was used^21-23^. We therefore determined whether the task-dependent modulation of connectivity by PMN-targeted stimulation occurred particularly for the PMN versus the ATN, using *a priori* defined regions of interest for each network^11^. Furthermore, we tested whether the positive influence of PMN-targeted TMS on the up-regulation of fMRI connectivity during memory retrieval would predict episodic memory improvement measured in an independent task. This allowed us to address the hypothesis that selective increases in fMRI connectivity during memory retrieval serves as an indicator of effective hippocampal network function that can be modulated by PMN-targeted TMS.

## Results

### Stimulation effects on task-dependent fMRI connectivity within hippocampal networks

We examined the task-dependent effects for PMN-targeted versus PFC-targeted control stimulation on network-wide interconnectivity (i.e., mean connectivity of each region to all other regions) in the PMN and the ATN using a linear mixed effects model with factors stimulation condition (stimulation versus sham), task (retrieval versus rest), stimulation location (PMN-targeted versus PFC-targeted) (see Methods: Data Analysis). The primary hypothesis was that there would be three-way interaction within the PMN among stimulation condition, task, and stimulation target, reflecting increased fMRI interconnectivity due to stimulation measured during autobiographical retrieval versus rest, selectively for PMN-targeted stimulation versus out-of-network PFC-targeted control stimulation. Indeed, stimulation effects on PMN interconnectivity were greater during retrieval versus rest following PMN-targeted stimulation relative to PFC-targeted stimulation (3-way interaction T(89)=2.80, p=0.006) (Figure 2A). This task-sensitive relative interconnectivity increase for PMN-targeted versus PFC-targeted stimulation was not found for the ATN (3-way interaction T(89)=1.48, p=0.14) (Figure 2A). Other main effects and interactions were nonsignificant in both networks (T<2, Supplementary Table 1).

**Figure 2:**
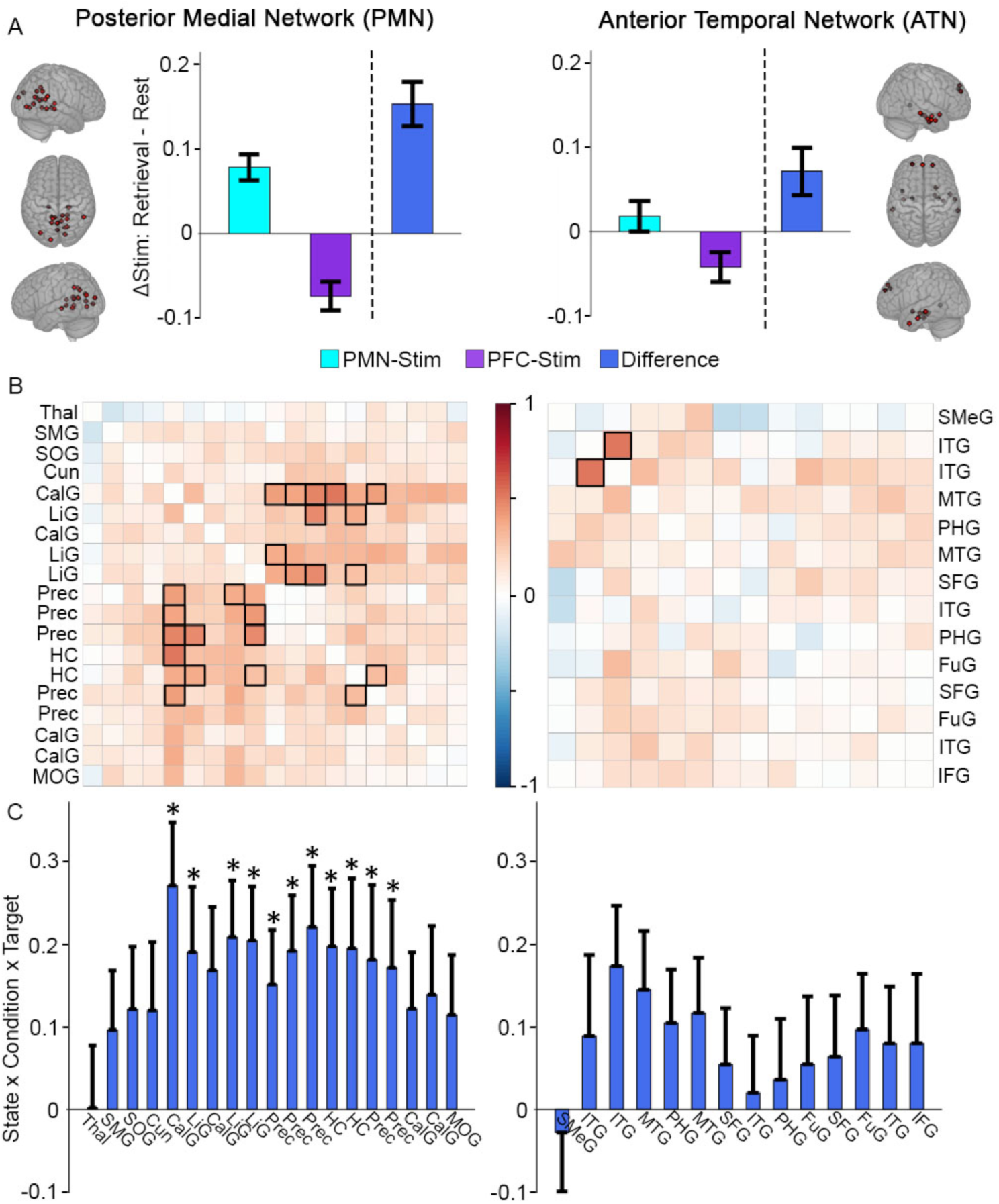
Greater memory task-dependent connectivity increases due to PMN-targeted versus PFC-targeted control stimulation for the PMN. (A) Stimulation effect on mean connectivity (all regions to all other regions) during memory retrieval relative to rest for the PMN (left) and the ATN (right). Individual bars are not marked for statistical significance as this would be redundant with values provided in the text. (B) The same effect of stimulation on connectivity between each pair of regions in the PMN (left) or ATN (right). Coloration represents the beta-weight of the three-way interaction effect, with significant cells shown in bold (FDR corrected p<0.05). (C) The same effect of stimulation on connectivity for each region in the PMN to all other PMN regions (left) and each region in the ATN to all other ATN regions (right). Error bars indicate subject-level standard error of the mean. * = FDR corrected p<0.05. Regions shown in Supplementary Figure 1A and region abbreviations are expanded in Supplementary Table 2.

We next identified regions that were driving these stimulation effects on network-wide task-dependent interconnectivity. Correlation matrices for the PMN and ATN were constructed, with the same linear mixed effects model used for each pair of regions as in the whole-network analysis to test the task-dependent effects on their connectivity following PMN-targeted versus PFC-targeted control stimulation. The pairwise connections showing significantly greater task-dependent stimulation effects during retrieval following PMN-targeted stimulation following correction for multiple comparisons (Figure 2B) included higher-order visual regions of the PMN, especially the calcarine and lingual gyri, with the hippocampus and precuneus. Notably, these were not the regions to which stimulation was directly applied (see Figure 1A), and hippocampal connectivity with these higher-order visual regions has been particularly implicated in episodic recollection^7,35^. We also computed average connectivity of each region to all other within-network regions to identify those regions that increased in task-dependent connectivity with the entire rest of the network. Within the PMN, the same regions showing pairwise task-dependent connectivity increases (hippocampus, precuenus, and calcarine and lingual gyri) also exhibited significant increases in task-dependent connectivity with the rest of the network, after correcting for multiple comparisons (Figure 2C). No ATN regions showed this effect.

### Increased whole-brain fMRI connectivity during memory retrieval

Following the targeted network analysis, the task-dependent effects of stimulation on fMRI connectivity were measured via whole-brain analyses^31,45,46^ of data from resting-state and retrieval task scans^31^ to allow for identification of regions without *a priori* designations of regions of interest. To validate the retrieval scan as an assay for memory-related connectivity and to show that our method for whole-brain fMRI connectivity measurement is sensitive to changes caused by memory retrieval, we first assessed the main effect of task (retrieval versus rest) independent from the factors stimulation type (stimulation versus sham) and stimulation location (PMN-targeted versus PFC-targeted) via voxel-wise linear mixed effects modeling (See Methods: Data Analysis). There was a main effect of task in the hippocampal-cortical network as well as regions typically associated with autobiographical memory, with greater connectivity during the retrieval task than during rest (Supplementary Figure 2). This replicates our previous findings using this analysis method and retrieval task^31^, and is consistent with many findings of fMRI connectivity increases due to similar memory retrieval tasks^47-50^. Therefore, our methods provide valid indicators of task-dependent changes in functional connectivity associated with memory.

### Exploratory whole-brain analysis of task-dependent effects of PMN-targeted stimulation

In order to evaluate the selectivity of effects of stimulation on the PMN and ATN, we next examined the task-dependent effects of stimulation using an exploratory, voxel-wise, whole-brain connectivity analysis approach^31,45,46^. We used the same linear mixed effects model as in the network analysis for the three-way interaction of condition, task, and stimulation location. Consistent with the targeted network analysis, there were many regions that exhibited significantly greater stimulation effects on whole-brain connectivity during retrieval versus rest following PMN-targeted stimulation relative to PFC-targeted control stimulation (Figure 3).

**Figure 3:**
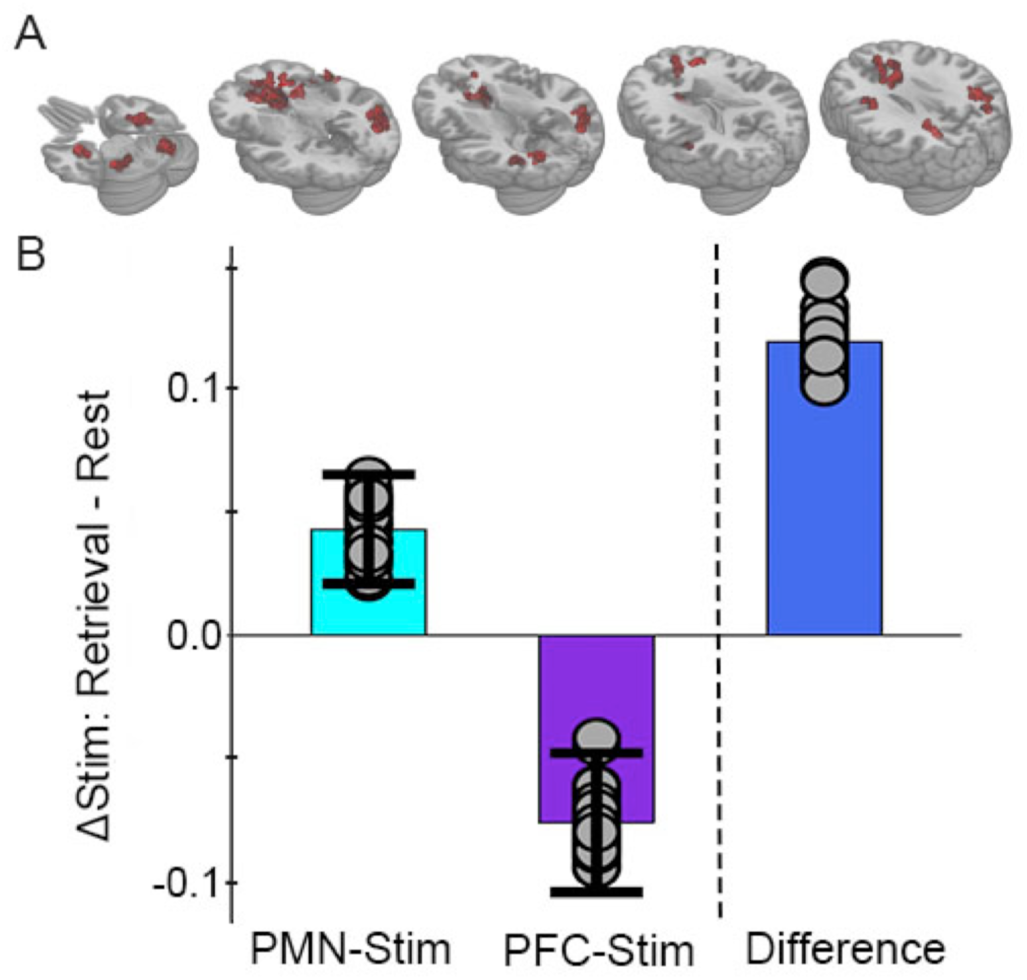
Greater memory task-dependent connectivity increases due to PMN-targeted versus prefrontal-control stimulation. (A) Regions showing a significant interaction between condition, task, and group, with red coloration indicating stimulation increased connectivity more during retrieval than during rest in the PMN-targeted group relative to the prefrontal-control group. All regions showed greater memory task-realted connectivity change following PMN-targeted stimulation. Comprehensive view shown in Supplementary Figure 3A. (B) Mean stimulation effect on connectivity during memory retrieval relative to rest for all supra-threshold regions. Error bars indicate subject-level standard error of the mean for supra-threshold regions. Points indicate the mean effect for each supra-threshold region. Note: Statistical values are not indicated, as this would be redundant with the statistical definition of these supra-threshold regions.

Because of the difference identified across stimulation groups, we next tested the interaction of stimulation condition (stimulation versus sham) and task (retrieval versus rest) using linear mixed effects models performed separately for each stimulation group (PMN-targeted or PFC-targeted; See Methods: Data Analysis). For the PMN-targeted stimulation group, there was significant interaction of stimulation by task in many regions particularly within the hippocampal-cortical network (Figure 4A). All but one area showed relatively greater connectivity for stimulation relative to sham during retrieval compared to rest, thereby demonstrating the hypothesized task-dependent increase in fMRI connectivity due to stimulation (Figure 4C).

**Figure 4:**
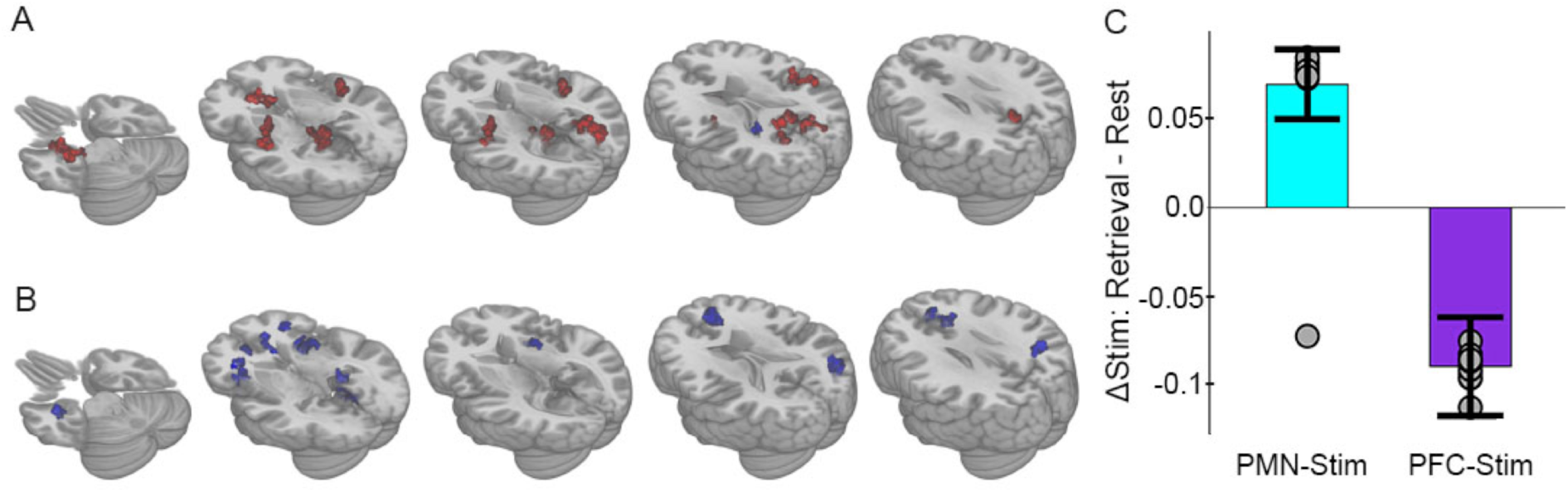
Selective effects of PMN-targeted stimulation on memory-related connectivity. (A) Regions showing significant interaction between stimulation condition and cognitive task following PMN-targeted stimulation, with red coloration indicating stimulation increased connectivity more during retrieval than during rest and blue coloration indicating the opposite effect. Comprehensive view shown in Supplementary Figure 3B-C. (B) Regions showing significant interaction between stimulation condition and cognitive task following PFC-targeted control stimulation. (C) Mean stimulation effect on connectivity during memory retrieval relative to rest for all supra-threshold regions. Error bars indicate subject-level standard error of the mean for all supra-threshold regions. Points indicate the mean effect for each supra-threshold region. Note: Statistical values are not indicated, as this would be redundant with the statistical definition of these supra-threshold regions.

The majority of regions (93.5%) demonstrating this interaction effect overlapped with the network that was sensitive to autobiographical retrieval (Supplementary Figure 2), indicating that task-dependent connectivity changes due to stimulation occurred for regions that contribute to memory retrieval. Furthermore, 61% of regions demonstrating the interaction effect were within *a priori* defined hippocampal-cortical network locations (8 PMN, 2 ATN, and 1 DMN, see Methods) with the remaining falling among anterior salience, sensorimotor, precuneus, and auditory networks that have been associated with autobiographical retrieval^47-50^. Thus, PMN-targeted stimulation had memory task-dependent effects primarily within hippocampal-cortical regions of *a priori* interest, with other areas showing this effect being involved generally in memory retrieval.

PFC-targeted control stimulation caused a nearly opposite pattern of fMRI connectivity change relative to PMN-targeted stimulation. As was the case for PMN-targeted stimulation, there was significant interaction of stimulation by task on connectivity (Figure 4B). However, all regions showed the opposite direction of connectivity changes relative to PMN-targeted stimulation, with greater connectivity during rest than during retrieval following PFC-targeted stimulation relative to sham (Figure 4C). Notably, the area of prefrontal cortex stimulated in the control condition is not part of the hippocampal-cortical network and participates in a variety of non-memory cognitive operations such as attention and maintenance of external awareness^51,52^. Stimulation of the control location may thus have caused a relative increase in these operations during rest and/or a disruption of memory-related processing during the retrieval task.

The locations of this interaction effect were consistent with results obtained from the three-way interaction model (Figure 3), which overlapped with 94.4% of the areas obtained via the PMN-targeted stimulation results and 86.7% of the PFC-targeted stimulation results. We then used a whole-brain analysis approach to thoroughly compare spatial distributions for all regions that contributed to connectivity effects in the group-level analyses (See Methods). Regions driving the task-dependent connectivity increases due to stimulation were categorized as belonging to PMN versus ATN^9,10^. As in the targeted analysis of effects on PMN and ATN (Figure 2), PMN-targeted stimulation produced memory task-dependent increases in connectivity in 31 PMN regions and only 5 ATN regions. In contrast, PFC-targeted stimulation produced memory task-dependent decreases in an evenly distributed set of PMN and ATN regions (12 PMN regions, 13 ATN regions). There was a significant difference in the relative distribution of task-dependent effects (irrespective of directionality) on PMN versus ATN regions in the PMN-targeted relative to PFC-targeted stimulation conditions (Yates-corrected X^2^(1) =8.56, p=0.003). Thus, the relatively selective effects of stimulation on PMN were consistent across the targeted analysis of PMN and ATN and whole-brain analyses.

### Increased memory task-dependent connectivity predicts episodic memory improvement

Episodic memory improvement was measured using an independent task (i.e., separate from the autobiographical retrieval and resting-state tasks used to assess task-dependent stimulation effects on fMRI connectedness). This task involved item recognition and context recollection (see Methods: Memory Task) (Figure 5A). Because context recollection is more heavily dependent on the PMN than item recognition^10,11^ and based on our previous findings^21-24^, we predicted that PMN-targeted stimulation would improve context recollection selectively^10^. Consistent with the previously reported results in 30 of the 32 subjects analyzed here^21^, PMN-targeted stimulation improved context recollection (T(15)=1.93, p=0.036) whereas PFC-targeted control stimulation did not (T(15)=-0.09, p=0.46), with significantly greater improvement for PMN-targeted stimulation relative to PFC-targeted stimulation (T(15)=1.88, p=0.04). Stimulation had no effect on item recognition for either stimulation location (PMN-targeted: T(15)=0.94, p=0.36; PFC-targeted: T(15)=-1.34, p=0.20).

**Figure 5:**
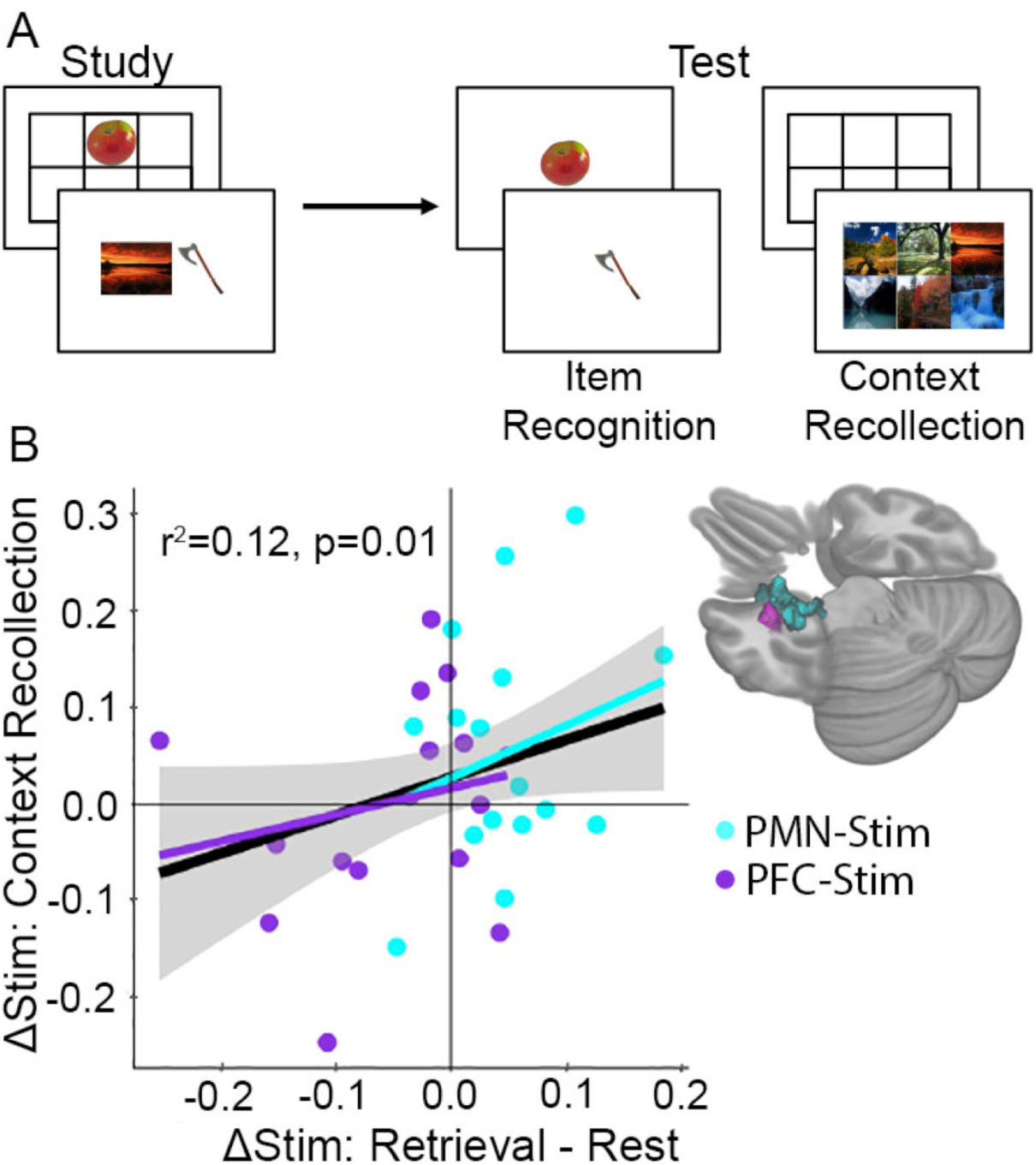
Increased memory-dependent connectivity predicts episodic memory improvement. (A) Episodic memory task design. Participants studied trial-unique objects paired with either scene or location contexts. After a delay, we assessed object recognition memory and contextual recollection memory. (B) Connectivity values were pooled across the MTL regions identified in each group showing an interaction between stimulation condition and cognitive task. Scatterplot shows the relationship between each subject’s change in retrieval task connectivity relative to rest and their change in context recollection performance following stimulation. Greater specificity of connectivity change to the memory task in the left MTL was associated with improvement in context recollection in each group independently (cyan and purple regression lines) as well as collectively across all 32 subjects (black regression line).

We next tested whether task-dependent increases in fMRI connectivity predicted memory improvement. Because of the critical role of the medial temporal lobe in memory^22,53^and on our previous findings showing left-lateralized effects of the same stimulation parameters on hippocampal fMRI activity^22,23^, we focused our analysis on left medial temporal lobe. The amount that PMN-targeted stimulation caused task-dependent increases in connectivity for retrieval versus rest was associated with greater improvement in context recollection memory (Robust regression F(15) = 6.64, r^2^=0.12, p=0.022). Although PFC-targeted stimulation was associated with net opposite task-dependent effects as PMN-targeted stimulation (i.e., greater connectivity increases for rest compared to retrieval), the same positive relationship was identified between memory task-dependent connectivity and context recollection as for PMN-targeted stimulation. That is, relative decreases in connectivity during retrieval than during rest predicted relative impairment in source recollection (Robust regression F(15)=11.14, r^2^=0.16, p=0.005). Based on the similarity of these effects, we pooled connectivity values across the two regions of interest and found that, irrespective of stimulation condition, the degree to which stimulation increased (rather than decreased) task-dependent connectivity during retrieval compared to rest predicted the amount of context recollection improvement (F(31) = 6.71, r^2^=0.12, p = 0.01) (Figure 5). Thus, memory task-dependent connectivity increases for the left medial temporal lobe is a robust indicator of episodic memory ability following stimulation.

## Discussion

These findings demonstrate memory task-dependent increases in the expression of fMRI connectivity changes caused by PMN-targeted stimulation. These effects were selective to *a priori* defined regions of the PMN relative to the ATN, which was confirmed via exploratory whole-brain analysis. Task-dependent connectivity increases in these regions were also specific to the PMN-targeted stimulation condition, relative to PFC-targeted control stimulation. Furthermore, retrieval-related connectivity increases in left medial temporal cortex and hippocampus due to stimulation predicted context recollection improvement measured in a separate task. Thus, enhancing hippocampal-cortical network connectivity during memory processing is functionally critical and is achievable via noninvasive stimulation. Furthermore, these task-dependent effects were measured ∼24 hours after stimulation, indicating that stimulation led to hippocampal-cortical network upregulation of connectivity during a subsequent memory task administered long after, and without any specific relationship to, stimulation delivery.

The PMN and ATN are functionally distinct components of the larger hippocampal-cortical network and are thought to differentially support memory processing related to context recollection versus item recognition, respectively^10,11^. Here, we report that increased connectivity due to stimulation was selective for PMN regions and for the autobiographical memory retrieval task. This selectivity is consistent with the role of the PMN in autobiographical memory retrieval^10^ as well as with the stimulation location, which was within the PMN^11^. These findings are consonant with our previous report that the same stimulation protocol increased stimulus-evoked activity during the encoding of item-context pairings selectively within the PMN^21^, and further supports hypothesized functional distinctions of the PMN and ATN.

Connectivity changes due to stimulation were measured during an autobiographical memory retrieval task versus rest, yet they predicted the effects of stimulation on memory ability measured using an independent item-context episodic memory test. Although autobiographical retrieval and episodic memory are superficially distinct, there is accumulating evidence that they are supported by similar cognitive operations and brain regions^49,54-56^. Our findings of a relationship of stimulation effects on connectivity during autobiographical retrieval with episodic memory performance underscores this similarity. Indeed, we have previously found that the PMN-targeted stimulation parameters used here improve a variety of episodic memory tasks of different formats, including face-word paired associates^22^, item-scene paired associates^21,23^, item-location associations^21,23^, and highly precise spatial recall^24^. Although we did not assess stimulation effects on the success of autobiographical memory retrieval, the current findings emphasize that stimulation targeting the PMN improves a variety of memory measures. Though these memory tasks are typically considered distinct (paired associate versus spatial recall, episodic versus autobiographical), they all have been associated with the PMN^6,47,49,54-56^ and therefore it is logical that they all respond to PMN-targeted stimulation.

Although resting-state fMRI connectivity has proven useful in characterizing the effects of stimulation on brain network function, including to understand memory improvements due to stimulation^19,20,22,33^, the current findings highlight one of the many limitations of resting-state fMRI for this purpose. That is, relationships between resting-state connectivity and network function are not well specified. Here, there was strong modulation of the effects of stimulation on connectivity based on whether it was measured during a memory retrieval task versus during rest, indicating that resting-state fMRI alone was an incomplete assay for stimulation effects. Furthermore, the functional relevance of stimulation for memory performance was predicted by the memory task-dependent change in connectivity, indicating that resting-state fMRI alone would not have identified functionally relevant effects of stimulation on connectivity.

It is likely that individuals engage in a variety of uncontrolled and typically unmeasured cognitive operations during resting-state fMRI and that these at least partially drive resting-state fMRI connectivity outcomes^51,52,57,58^. Our finding that stimulation effects on connectivity are particularly strong during a memory retrieval task is especially problematic for resting-stat fMRI as an outcome for memory interventions given that subjects frequently and variably retrieve memories during resting-state fMRI^51,52,57,58^. Variable effects of stimulation on fMRI connectivity at rest would therefore be expected based on the content and quality of memory retrieval subjects experience during scanning. The current experiment accounted for these challenges by giving an explicit retrieval task during connectivity measurement, which permitted differentiation of the PMN-targeted versus PFC-targeted stimulation conditions and identified functionally relevant effects of stimulation on memory performance.

To summarize, our findings suggest that PMN-targeted brain stimulation increases activity coupling among PMN regions when these regions are engaged by memory processing demands, rather than nonspecifically engaged during rest. Furthermore, task-dependent connectivity increases in the medial temporal lobe predicted improvement in a separate memory task. Thus, memory enhancement by brain stimulation relies on dynamic (i.e., activity-dependent) rather than static changes in connectivity among select portions of the hippocampal-cortical network, thereby reflecting a form of effective rather than functional connectivity^59^. The relationship between task-dependent connectivity and memory improvement was robust for the left hippocampus and surrounding medial temporal cortex, consistent with its established role in memory encoding and retrieval^4,5,10^. This finding is important given that nonspecific increases in connectivity could be detrimental to memory as well as other cognitive abilities, as this would entail less-selective participation of the hippocampal-cortical network in memory^18,29-36^. Stimulation-based interventions for memory disorders should therefore strive to achieve memory task-dependent functional engagement, as identified here using PMN-targeted noninvasive stimulation.

## Methods

### Participants

Thirty-two adults participated in the experiment (22 females, mean age = 25.6 years, range = 18-34). Data from two additional participants were collected but discarded due to excessive motion (see below). All conditions of interest were fully counterbalanced in the final sample contributing data to analyses. All participants had normal or corrected-to-normal vision and did not report a history of neurological or psychiatric disorders or current drug use. Participants were eligible for MRI and TMS procedures according to standard MRI and TMS safety-screening questionnaires. Eligibility contraindications were evaluated by a neurologist (S.V.) Memory performance data from 30/32 subjects contributing to the analysis of the relationship between connectivity and memory (see below) has been previously reported^21^. The Institutional Review Board at Northwestern University provided approval for this study. All participants gave written, informed consent and were paid for their participation.

### Experiment Design

Participants completed a 2-week experiment involving one week of full-intensity stimulation and one week of sham stimulation (Figure 1A) to a target in either a posterior medial network target in the left lateral parietal lobe (PMN-targeted stimulation, N=16) or a control site in the left lateral prefrontal cortex (PFC-targeted stimulation, N=16). The order of these two weeks was counterbalanced, and the first day of each week was separated by a delay of at least 4 weeks (mean interval=11.52 weeks, range=4.71–37.14 weeks). About 2 hours before receiving stimulation on the first day of each week and ∼24 hours after five consecutive daily stimulation sessions (mean delay=23.3, SD=2.50 hours from the final stimulation session), participants completed a resting-state scan, an autobiographical memory retrieval task scan, and a task-based fMRI memory paradigm (Figure 1B, Figure 4A). Task-associated behavioral and fMRI data have been reported elsewhere for 30 of the 32 subjects^21^, with two subjects replaced due to excessive motion during resting-state scans to achieve the full sample reported here (N=32). The present analyses focus on post-stimulation versus post-sham comparisons, as these have been shown to isolate the effects of stimulation^21,23,24^.

### Resting-State and Autobiographical Memory Retrieval Task Scans

The resting-state and the autobiographical memory retrieval task scans used the same EPI sequence, which each lasted 5.5 minutes. During the resting-state scan, a fixation cross was presented continuously, and participants were instructed to remain awake with eyes open and fixated on the cross. During the retrieval task scan, participants were shown text prompts for common life events, such as “A graduation you attended” or “A vacation you enjoyed,” and were instructed to vividly recall these events (Figure 1C)^31^. Each prompt was shown for 20-s with a 10-s inter-prompt interval, and participants were told to imagine the event for the full 30-s period. Ten event prompts were shown consecutively during the scan. There were six sets of prompts each including distinct events. Each subject received a different version at each assessment, with order counterbalanced across subjects receiving full stimulation and sham stimulation during the first week.

### Imaging Acquisition and Processing

Participants were scanned using a Siemens MAGNETOM Prisma scanner with a 64-channel head/neck coil. The resting-state and retrieval task scans were both acquired using a T2* weighted echo planar imaging (EPI) sequence (TR = 555 ms, TE = 22 ms, multiband factor=8, isotropic voxel size = 2×2×2mm, FOV = 208×192 mm, flip angle = 47°, volumes = 550). Structural imaging was acquired using MPRAGE T1-weighted scans (TR = 2170 ms, TE = 1.69 ms, voxel size = 1 mm^3^, FOV = 25.6 cm, flip angle = 7°, 176 sagittal slices).

Data were preprocessed using AFNI (Version AFNI_16.1.15), with the same processing steps used for retrieval task and resting-state scans. The first five EPI volumes were removed to avoid intensity normalization artifact. AFNI’s *3dDespike* was used to remove large transient volumes. The remaining volumes were slice-time and multiband corrected. Six estimated motion parameters (x, y, z, roll, pitch, and yaw) were estimated using 3dvolreg. Data from two participants were excluded for high motion (17% and 13% of TRs censored at FD threshold 0.3mm; <5% censored for all included subjects). For included participants, pairwise comparisons were made for post-stim, post-sham, retrieval task, and resting-state scan framewise displacement (FD) and temporal signal to noise ratio (tSNR) in the PMN-targeted and PFC-targeted control stimulation groups. All FD comparisons were nonsignificant (p > 0.05), however, marginal differences in temporal signal to noise ratio (tSNR) were found between post-stimulation and post-sham retrieval scans in the PMN-targeted stimulation group (T(15)=-2.18, p=0.05), thus all statistical analyses used tSNR as a covariate of no interest. Volumes were co-registered to the anatomical scan and then transformed into standardized Talairach and Tournoux space (TT_N27 atlas). Images were then smoothed with an isotropic 4.0 mm full-width half-maximum Gaussian kernel and masked using AFNI’s *3dAutomask*. The six motion parameters estimated earlier were then regressed out of the timeseries using AFNI’s *3dDeconvolve*, after which a bandpass filter was applied (0.01-0.1 Hz) using AFNI’s *3dTproject*. A group mask was created by merging masks of all subjects using AFNI’s *3dmerge* and including only voxels that were present in the final preprocessed datasets in all subjects.

### Identification of stimulation locations

Individualized left lateral parietal (PMN-targeted) or left prefrontal (PFC-targeted control) stimulation locations were determined based on high resting-state fMRI connectivity with a left hippocampal seed using a procedure previously described^21,22,24^. Briefly, resting state data from the first visit was used to select a hippocampal volume of interest for each participant by identifying a voxel in the body of the left hippocampus closest to MNI [−29, −25, −13] (mean distance=2.51mm, range=0.00–6.71mm) for which fMRI connectivity was maximal to contralateral hippocampus. This location was used for seed-based connectivity analysis (AFNI’s *InstaCorr*) using a seed radius of 2 mm.

For the PMN-targeted stimulation group, the stimulation location was selected as the peak connectivity cluster within left lateral parietal cortex, within an anatomical mask of angular and supramarginal gyri and inferior parietal lobule close to MNI [−47, −68, 36] (mean distance=10.2mm, range=0.0-35.8mm from this coordinate; Figure 1B). For the PFC-targeted stimulation group, the stimulation location was selected as the peak connectivity cluster within a functional mask of left dorsolateral prefrontal cortex close to MNI [-23, 40, 43] (mean distance=10.5mm, range=0.0-19.9mm; Figure 1B). This mask was generated by Neurosynth as meta-analytic co-activation with the left hippocampus (MNI [−29, −25, −13]). The stimulation target was transformed for each participant to original space for anatomically guided stimulation. The same stimulation location was used for each subject for both stimulation and sham weeks.

### Transcranial Magnetic Stimulation (TMS)

The MagPro X100 system with a MagPro Cool-B65 liquid-cooled butterfly coil was used (MagVenture A/S, Farum, Denmark). A frameless stereotactic system (Localite GmbH, St. Augustin, Germany) used individual MRIs for anatomical targeting of stimulation and for recording coil locations relative to the brain for each TMS pulse. Resting motor threshold (MT) was determined visually based on the minimum stimulator output required to generate a contraction of the *abductor pollicis brevis* for 5 out of 10 consecutive single pulses. Repetitive TMS was planned at 100% MT for each day of the stimulation week and at 10% MT for each day of the sham week, although these values were lowered due to discomfort for 5 subjects in the PMN-targeted stimulation group (to 95%, 90%, 82%, 80%, and 72% MT) and for 3 subjects in the PFC-targeted group (to 89%, 83%, and 74% MT). The final mean stimulator output intensity for full stimulation in the PMN-targeted group was 52.5 (SD=8.5) and 51.2 (SD=7.7) for the PFC-targeted group (T(31)=0.22, p=0.83). The mean output for sham was 5.42 (SD=0.90) in the PMN-targeted group and 5.19 (SD=0.66) for the PFC-targeted group. Each daily TMS session consisted of 40 consecutive trains of 20-Hz pulses for 2 seconds followed by 28 seconds of no stimulation (1600 pulses per session, 20 minutes total).

### fMRI Data Analysis

Two components of the hippocampal-cortical network, the PMN and the ATN, were of particular interest, as the stimulation location was within the PMN and we have previously shown that the effects of this same stimulation protocol on task-based fMRI activity during memory encoding are greater for regions within the PMN network than the ATN^21^. Regions in the PMN and ATN were defined *a priori* based on previous studies of fMRI connectivity with parahippocampal and perirhinal cortex, respectively^11,60^. Network regions were 6-mm-radius spheres centered on the peak coordinates of each network location (Supplementary Figure 1A). The spatially averaged time series for each region was extracted from each subject’s post-stimulation and post-sham retrieval and resting-state scans and used to construct a correlation matrix for each state and condition (R package *corrplot*, RStudio 1.1.453). Network connectivity was first assessed as the average of all pairwise correlations within a network (Figure 2A). This was compared among conditions using a linear mixed effects model (R packages *lme4* and *lmerTest*) with the factors stimulation condition (stimulation/sham), cognitive task (retrieval/rest), stimulation group (PMN-targeted/PFC-targeted), their three-way interaction, and a control tSNR term (see Image Acquisition and Processing). To identify which connections contributed to the average network effects, we then used the same three-way interaction model to identify effects on connectivity for each pair of regions (Figure 2B). Significant links were defined by a two-tailed pair-wise f-value threshold of p<0.05 after FDR correction. Finally, we identified which regions showed the greatest task-dependent stimulation effects by averaging each region’s correlation with all other network regions and comparing these values with the three-way interaction model (Figure 2C). Significant regions were defined by a two-tailed voxel-wise f-value threshold of p<0.05 after FDR correction.

Next, we expanded our analysis to examine the effects of stimulation on whole-brain fMRI connectivity. Following our previous study examining retrieval task and resting-state differences^31^, we used a whole-brain global connectedness analysis to identify the differential effects of stimulation on resting-state and retrieval fMRI connectivity. Global fMRI connectedness maps were created for each participant’s post-stimulation and post-sham rest and retrieval fMRI scans separately^31,45,46^. The correlation of each voxel’s timeseries was computed against every other voxel within the brain mask and the mean correlation with all other voxels was stored back into the voxel (AFNI’s *3dTcorrMap*), giving a measure of how correlated each voxel is with all other voxels throughout the entire brain. These global fMRI connectedness maps were transformed using Fisher’s z to create normally distributed values. This method is data-driven yet conservative, as any significant correlations must survive being washed out by weaker or opposite-direction correlations in other voxels throughout the entire brain. Regions identified in using this data-driven approach were then codified based on membership to well-characterized functional networks to aid interpretation.

Mean connectivity was compared among conditions using linear mixed effects models using AFNI’s *3dLME*. We first used the same model as in the PMN and ATN ROI analyses, which included the factors stimulation condition (stimulation/sham), cognitive task (retrieval/rest), stimulation group (PMN-targeted/PFC-targeted), their three-way interaction, and a control tSNR term (see Image Acquisition and Processing) (Figure 3). We then created two models which independently examined PMN-targeted and PFC-targeted group results with the factors stimulation condition, cognitive task, their interaction, and tSNR (Figure 4). A grey matter mask was then applied to exclude any regions falling in white matter, created by averaging the MPRAGE scans of all 32 subjects, which was then used to create the grey matter mask using AFNI’s *3dSeg*. Significant clusters were defined by a two-tailed voxel-wise f-value threshold of p<0.05, which is typical for fMRI connectedness given that experimental effects of subsets of voxels are averaged with null effects for the majority of the brain^31,45,46^ and that two-tailed testing avoids inflation of false positive results present in the majority of neuroimaging experiments using one-tailed testing^61-63^. We controlled false positives by computing a threshold for the minimum number of contiguous supra-threshold voxels using permutation testing. Permutation testing was conducted by running the three-factor model 1000 times with random flipping of factor labels. A probability distribution of cluster sizes was generated across all permutations for each factor using the two-tailed f-value threshold of p<0.05 from our primary analysis. Cluster size cutoffs were then defined as the size with only a 5% probability of finding any cluster that size or larger given random factor label assignment. This identified a threshold of 38 voxels for the stimulation main effect, 36 voxels for the cognitive task main effect, and 36 voxels for the three-way interaction effect. We applied the most stringent of these thresholds, 38 voxels, to all effects to identify significant results. On average, supra-threshold clusters were 2.2 times as large as this threshold (83.3 voxels).

Based on previous work^21,22^, we had *a priori* hypotheses that changes in the medial temporal lobe and the episodic memory network^10^ would be particularly related to changes in memory performance following PMN-targeted stimulation. We therefore characterized the network allegiance of the global correlation regions and their “drivers”, the locations which showed changes in connectivity with the global correlation regions and thus were contributing to the effect. To visualize the drivers, regions identified by the interaction effect in the independent group models (PMN-targeted or PFC-targeted) were used as seeds for voxel-wise whole-brain fMRI correlation analysis. The spatially averaged time series for each region was extracted from each subject’s post-stimulation and post-sham retrieval and resting-state scans and correlated with the time series of every other voxel in the brain mask. Seed-based correlation maps across subjects were then compared using the same two-factor linear model as in the global correlation step (AFNI’s *3dLME*). Significant clusters were defined by a voxel-wise f-value threshold of p<0.001 two-tailed and a cluster extent threshold of 20 contiguous voxels in the interaction effect. This is a highly conservative threshold for all recent thresholding recommendations^61-63^. We again examined the relationship between these effects and the PMN and ATN ROI effects first identified. Clusters from the whole-brain analysis which overlapped with one of the network regions were considered as clusters within that network. Network allegiance of all remaining clusters was identified using an atlas of 14 resting-state functional networks identified in Shirer, et al., 2012^64^ (Supplementary Figure 1B).

### Episodic memory assessment and analysis of its relationship to fMRI connectivity

The memory task completed at each assessment (post-stimulation and post-sham) involved study-test blocks and used two stimulus formats (Figure 5A), as described in Kim et al. 2018. For each block, participants studied 42 trial-unique objects either paired with one of six scenes or displayed in one of six locations on a grid, and then memory was tested after a 2-minute delay. During test blocks, half of trials were old (studied) objects and half of trials were new (unstudied) objects. Participants first categorized object as “old” or “new” and simultaneously rated confidence as “certain” or “uncertain” using four response options, providing a measure of item recognition memory. All studied objects were then tested for contextual recollection memory, whereby participants selected the scene or the location associated with the object during the study phase. Behavioral data have been reported previously^21^ for 30 of the 32 subjects included in the present study (i.e., there are only two new memory-task datasets for the current experiment, for the two subjects that were needed to replace those previously excluded due to poor resting-state fMRI data quality), showing effects of stimulation specifically on context recollection. Therefore, the effects of stimulation on memory performance reported below are largely redundant with the Kim et al. 2018 report and not considered novel evidence for stimulation effects on memory accuracy. Instead, these data are included here for the analysis of the relationship between stimulation effects on fMRI connectivity and context recollection accuracy.

At each assessment, item recognition memory was assessed as the proportion of total trials that were correctly recognized as old (hits) or new (correct rejections). Contextual recollection accuracy was assessed as the proportion of correct responses to the original scene or location context (one of six options) given that the object was correctly recognized with high confidence (“certain” response). Based on previous work with these data^21^ and *a priori* hypotheses of memory improvement due to parietal stimulation, as already reported in Kim et al. 2018, directional (one-tailed) paired t-tests were used to verify that stimulation effects on context recollection remained using the two replacement subjects in the current analysis. To reiterate, the goal of including these behavioral data was not to demonstrate the effects of stimulation on memory performance, which have already been reported for 30/32 subjects in the current report, but rather in order to assess the relationships between effects of stimulation on fMRI connectivity and on memory performance.

We identified associations between changes in memory performance and task-dependent stimulation effects on global correlativity for each group (individual group model interaction effects) (Figure 5). We focused on the *a priori* hypothesis that the left medial temporal lobe would show the greatest effects based on previous findings of left-lateralized effects of PMN-targeted stimulation on hippocampal fMRI activity^21,23^ and on the general importance of medial temporal lobe for memory^4,5^. For each of the interaction effect clusters within the left medial temporal lobe for both PMN-targeted and PFC-targeted stimulation conditions, a robust linear model was built regressing the interaction effect for that cluster (retrieval task stimulation effect minus resting-state stimulation effect) and tSNR onto the stimulation effect on memory performance (post-stimulation minus post-sham) (R packages *MASS* and *sfsmisc*, RStudio 1.1.453).

## Acknowledgements

This project was funded by R01MH106512 from the National Institute of Mental Health. The content is solely the responsibility of the authors and does not necessarily represent the official views of the National Institutes of Health. The authors would like to thank Stephen VanHaerents for his assistance with TMS safety screening, Sungshin Kim for his contributions to the episodic memory task, and Jonathan O’Neil, Robert Palumbo, Elise Gagnon, Melissa McSweeney, and Melissa Gunlogson for their roles in data collection. This research was supported in part through the computational resources and staff contributions provided for the Quest high-performance computing facility at Northwestern University, which is jointly supported by the Office of the Provost, the Office for Research, and Northwestern University Information Technology. Neuroimaging was performed at the Northwestern University Center for Translational Imaging, supported by Northwestern University Department of Radiology.

## Competing Interests

The authors have no competing interests to disclose.

## Author Contributions

J.L.V. designed the experiment. K.N.W., M.S.H., and A.S.N. assisted with data collection and ensured data quality. K.N.W. conducted analyses for all figures and, with J.L.V., interpreted results. K.N.W. wrote, and K.N.W., J.L.V., A.S.N, and M.S.H edited, the manuscript.

