## Supplementary Materials for "Increased task-dependent functional connectivity by stimulation of the hippocampal network predicts memory enhancement"

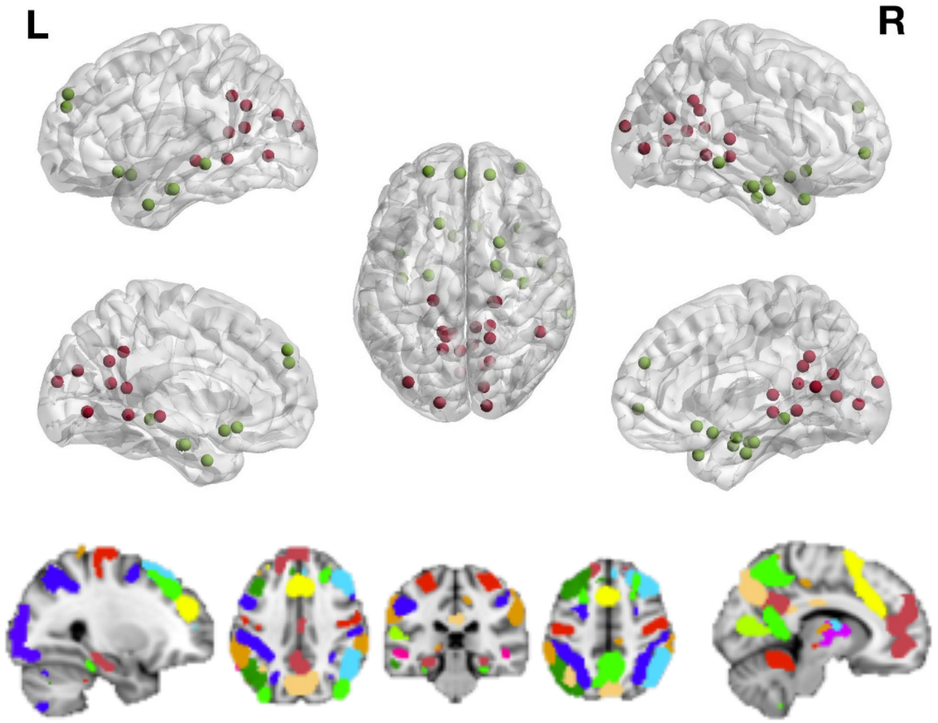


**Supplementary Figure 1: Network regions of interest. (A)** Posterior medial network (PMN) and anterior temporal network (ATN) regions of interest (Ranganath & Ritchey, 2012): PMN (red), ATN (green). **(B)** Depiction of 14 functional network regions of interest (Shirer, et al., 2012): Anterior salience (yellow), auditory (bright green), basal ganglia (purple), dorsal default mode (maroon), higher visual (blue), language (pink), left executive control (green), sensorimotor (red), posterior salience (orange), precuneus (tan), primary visual (light green), right executive control (light blue), ventral default mode (light green), visuospatial (blue).

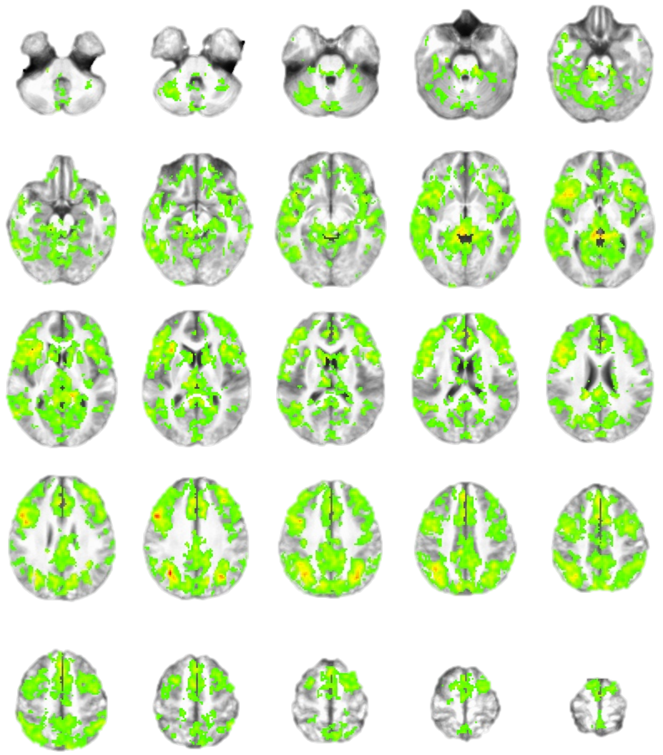

**Supplementary Figure 2: Increased fMRI connectivity during memory retrieval.** Regions showing a significant main effect of demand independent from stimulation condition or stimulation group. All showed greater connectivity during the retrieval demand relative to rest. Left=Left.

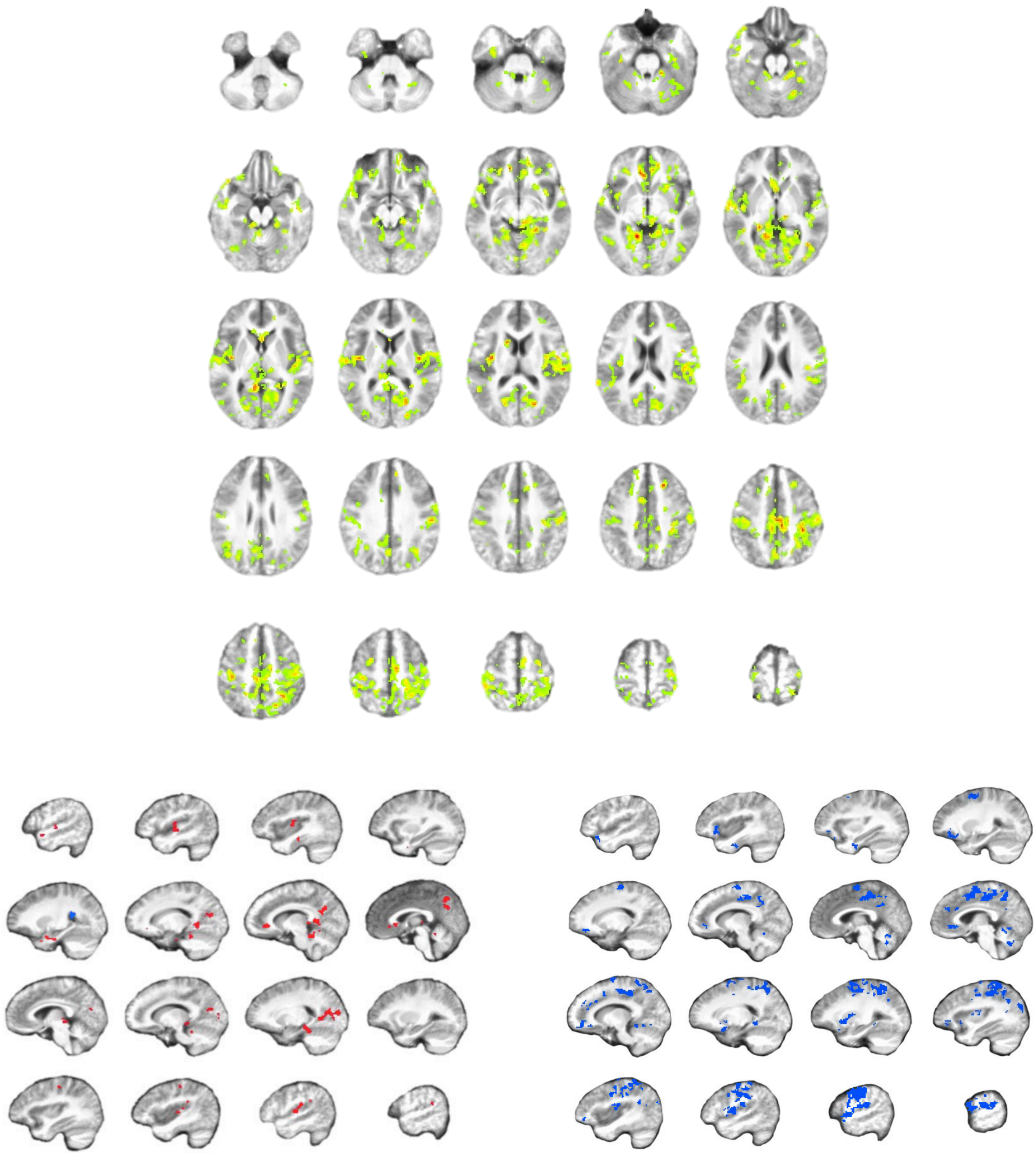

**Supplementary Figure 3: Selective effects of stimulation on memory-specific connectivity.** **(A)** All regions showing a significant three-way interaction between condition, demand, and group. **(B-C)** All regions showing a significant interaction between condition and demand following **(B)** HCN-targeted stimulation or **(C)** out-of-network control stimulation. Note: These are the same contrasts shown in Figures 3-4, but with all slices shown for a comprehensive view.

**Supplementary Table 1: PMN and ATN effect tables**

| PMN | | | | | |
|---|---|---|---|---|---|
| **Effect** | **Beta Estimate** | **Std. Error** | **df** | **t** | **p** |
| (Intercept) | 0.768 | 0.089 | 112.100 | 8.627 | 0.000 |
| State | 0.054 | 0.029 | 91.010 | 1.846 | 0.068 |
| Condition | 0.035 | 0.029 | 89.610 | 1.197 | 0.234 |
| Target | 0.016 | 0.040 | 70.880 | 0.399 | 0.691 |
| State x Condition | -0.063 | 0.041 | 89.730 | -1.553 | 0.124 |
| State x Target | -0.066 | 0.041 | 89.550 | -1.622 | 0.108 |
| Condition x Target | -0.062 | 0.041 | 90.180 | -1.505 | 0.136 |
| State x Condition x Target | 0.161 | 0.058 | 89.600 | 2.800 | 0.006* |
| **ATN** | | | | | |
| **Effect** | **Beta Estimate** | **Std. Error** | **df** | **t** | **p** |
| (Intercept) | 0.522 | 0.077 | 113.000 | 6.810 | 0.000 |
| State | 0.012 | 0.025 | 90.770 | 0.488 | 0.627 |
| Condition | 0.037 | 0.025 | 89.390 | 1.497 | 0.138 |
| Target | -0.013 | 0.035 | 68.670 | -0.372 | 0.711 |
| State x Condition | -0.041 | 0.035 | 89.510 | -1.176 | 0.243 |
| State x Target | -0.041 | 0.035 | 89.330 | -1.175 | 0.243 |
| Condition x Target | -0.019 | 0.035 | 89.960 | -0.539 | 0.592 |
| State x Condition x Target | 0.073 | 0.049 | 89.380 | 1.482 | 0.142 |

Effect tables for the linear mixed models used to identify the demand-specific effects for PMN-targeted versus PFC-targeted control stimulation (Figure 1).

**Supplementary Table 2: PMN and ATN ROI abbreviations**

| PMN | | | | |
|---|---|---|---|---|
| | **Region** | **x** | **y** | **z** |
| Thal | Thalamus | -21 | 28 | 8 |
| SMG | Supramarginal Gyrus | -49 | 47 | 26 |
| SOG | Superior Occipital Gyrus | 15 | 92 | 17 |
| Cun | Cuneus | -17 | 66 | 21 |
| CalG | Calcarine Gyrus | -13 | 68 | 7 |
| LiG | Lingual Gyrus | 13 | 47 | -4 |
| CalG | Calcarine Gyrus | -13 | 82 | 2 |
| LiG | Lingual Gyrus | 2 | 73 | -2 |
| LiG | Lingual Gyrus | -17 | 43 | -2 |
| Prec | Precuneus | -17 | 51 | 32 |
| Prec | Precuneus | 2 | 59 | 30 |
| Prec | Precuneus | 11 | 50 | 36 |
| HC | Hippocampus | 19 | 27 | -5 |
| HC | Hippocampus | -19 | 28 | -3 |
| Prec | Precuneus | 7 | 48 | 13 |
| Prec | Precuneus | -8 | 44 | 15 |
| CalG | Calcarine Gyrus | 13 | 58 | 16 |
| CalG | Calcarine Gyrus | -6 | 55 | 13 |
| MOG | Middle Occipital Gyrus | 35 | 79 | 24 |
| **ATN** | | | | |
| | **Region** | **x** | **y** | **z** |
| SMeG | Superior Medial Gyrus | 2 | -54 | 37 |
| ITG | Inferior Temporal Gyrus | 39 | 13 | -23 |
| ITG | Inferior Temporal Gyrus | -60 | 13 | -22 |
| MTG | Middle Temporal Gyrus | 60 | 33 | -7 |
| PHG | Parahippocampal Gyrus | -21 | 5 | -19 |
| MTG | Middle Temporal Gyrus | -66 | 35 | -7 |
| SFG | Superior Frontal Gyrus | 22 | -55 | 29 |
| ITG | Inferior Temporal Gyrus | 41 | -3 | -33 |
| PHG | Parahippocampal Gyrus | 22 | 11 | -23 |
| FuG | Fusiform Gyrus | -28 | 11 | -29 |
| SFG | Superior Frontal Gyrus | -17 | -53 | 28 |
| FuG | Fusiform Gyrus | -38 | 16 | -22 |
| ITG | Inferior Temporal Gyrus | -51 | 3 | -24 |
| IFG | Inferior Frontal Gyrus | -23 | -10 | -16 |

Expanded names for PMN and ATN region abbreviations used in Figure 2 and corresponding MNI coordinates.

**Supplementary Table 3: Findings of the task-dependent connectivity analysis and their corresponding drivers for PMN-targeted stimulation.**

| Cluster Peak (RAI) | | | |
|---|---|---|---|
| x | y | z | Region |
| **-21** | **65** | **14** | **Calcarine Gyrus** |
| -9 | 59 | 54 | Precuneus |
| 49 | -9 | -8 | Superior Temporal Gyrus/Temporal Pole |
| -1 | -37 | 36 | Superior Medial Gyrus |
| -49 | -15 | 28 | Inferior Frontal Gyrus |
| -29 | 3 | 58 | Superior Frontal Gyrus |
| -21 | -23 | 42 | Middle Frontal Gyrus |
| -55 | 49 | 2 | Middle Temporal Gyrus |
| -7 | -17 | 36 | Middle Cingulate Gyrus |
| **9** | **45** | **10** | **PCC/Precuneus** |
| -37 | 65 | 38 | Angular Gyrus |
| -45 | -37 | 2 | Inferior Frontal Gyrus |
| 47 | -7 | 32 | Precentral Gyrus |
| -43 | -7 | 46 | Middle Frontal Gyrus |
| **39** | **9** | **12** | **Insula** |
| 53 | -11 | -2 | Superior Temporal Gyrus/Temporal Pole |
| -7 | -35 | 32 | Anterior Cingulate Cortex |
| -59 | 33 | -12 | Inferior Temporal Gyrus |
| 47 | 55 | -6 | Inferior Temporal Gyrus |
| -35 | -29 | 14 | Inferior Frontal Gyrus |
| -17 | 57 | 12 | Calcarine Gyrus |
| -19 | 65 | 48 | Superior Parietal Lobule |
| 17 | 59 | 10 | Calcarine Gyrus |
| -5 | 15 | 54 | SMA |
| 43 | 19 | 44 | Postcentral Gyrus |
| -7 | -47 | -8 | Rectal Gyrus |
| **-19** | **35** | **-16** | **Cerebellum** |
| 11 | 73 | 24 | Cuneus |
| -15 | 53 | -8 | Lingual Gyrus |
| -23 | 77 | -18 | Cerebellum |
| -39 | 61 | -22 | Cerebellum |
| 47 | 47 | 14 | Middle Temporal Gyrus |
| 39 | -3 | 40 | Precentral Gyrus |
| -31 | 63 | -36 | Cerebellum |
| 17 | 57 | 38 | Superior Parietal Lobule |
| 55 | 43 | 42 | Inferior Parietal Lobule |
| 33 | -33 | 4 | Inferior Frontal Gyrus |

| | | | |
|---|---|---|---|
| 63 | 37 | 14 | Superior Temporal Gyrus |
| -3 | -21 | 28 | Anterior Cingulate Cortex |
| 17 | -31 | 32 | Superior Frontal Gyrus |
| 43 | -1 | 40 | Precentral Gyrus |
| -37 | 33 | 56 | Postcentral Gyrus |
| **9** | **-31** | **0** | **ACC** |
| -49 | -17 | 8 | Inferior Frontal Gyrus |
| 7 | -27 | 28 | Anterior Cingulate Cortex |
| 39 | -37 | 12 | Inferior Frontal Gyrus |
| 57 | 53 | 18 | Middle Temporal Gyrus |
| 27 | -47 | 4 | Middle Orbital Gyrus |
| 41 | 13 | 54 | Precentral Gyrus |
| 9 | 73 | -34 | Cerebellum |
| 1 | -19 | 48 | Superior Medial Gyrus |
| -1 | -47 | 28 | Superior Medial Gyrus |
| -19 | -9 | 4 | Putamen |
| 51 | -3 | 24 | Inferior Frontal Gyrus |
| 47 | 33 | 22 | Superior Temporal Gyrus |
| -3 | 29 | 26 | Middle Cingulate Gyrus |
| 23 | 59 | 16 | Calcarine Gyrus |
| -19 | -27 | 32 | Superior Frontal Gyrus |
| 15 | -11 | 60 | Superior Frontal Gyrus |
| -43 | 21 | 44 | Postcentral Gyrus |
| 17 | 9 | 62 | SMA |
| -5 | 71 | 12 | Calcarine Gyrus |
| **9** | **39** | **-16** | **Cerebellum** |
| 17 | 51 | 2 | Lingual Gyrus |
| -11 | 57 | -4 | Lingual Gyrus |
| 7 | 81 | 14 | Cuneus |
| 11 | 73 | -4 | Lingual Gyrus |
| -31 | 47 | 44 | Inferior Parietal Lobule |
| -47 | 39 | 52 | Superior Parietal Lobule |
| 47 | -15 | 10 | Inferior Frontal Gyrus |
| 49 | 57 | 30 | Angular Gyrus |
| **21** | **15** | **-22** | **MTL** |
| -17 | 45 | 32 | Middle Cingulate Gyrus |
| 41 | 1 | 44 | Precentral Gyrus |
| 53 | -5 | 22 | Inferior Frontal Gyrus |
| 47 | 45 | 6 | Middle Temporal Gyrus |
| 1 | 1 | 58 | SMA |
| **-47** | **13** | **8** | **Heschls Gyrus** |
| -7 | 45 | 10 | Precuneus |

| | | | |
|---|---|---|---|
| 5 | 51 | 56 | Precuneus |
| 43 | 55 | 0 | Middle Temporal Gyrus |
| 15 | 63 | -8 | Lingual Gyrus |
| 5 | -3 | 44 | SMA |
| -5 | -21 | 42 | Superior Medial Gyrus |
| 27 | 13 | 52 | Precentral Gyrus |
| -21 | 5 | 56 | Superior Frontal Gyrus |
| 51 | -11 | 38 | Middle Frontal Gyrus |
| -37 | 75 | -8 | Inferior Occipital Gyrus |
| 55 | -13 | 2 | Superior Temporal Gyrus/Temporal Pole |
| 51 | 67 | -6 | Inferior Occipital Gyrus |
| **19** | **41** | **16** | **Caudate** |
| -25 | 65 | -22 | Cerebellum |
| 7 | 49 | 58 | Precuneus |
| 49 | 39 | 4 | Middle Temporal Gyrus |
| 41 | 21 | -4 | Superior Temporal Gyrus |
| 51 | 13 | 4 | Superior Temporal Gyrus |
| **-35** | **19** | **46** | **Precentral Gyrus** |
| 3 | 29 | -34 | Brainstem |
| 31 | 67 | -4 | Inferior Occipital Gyrus |
| -25 | 79 | -16 | Fusiform Gyrus |
| 5 | 93 | 10 | Cuneus |
| 1 | 61 | 36 | Precuneus |
| 23 | -9 | 46 | Middle Frontal Gyrus |
| 1 | 53 | 56 | Precuneus |
| **-49** | **23** | **18** | **Insula** |
| 43 | 49 | 44 | Inferior Parietal Lobule |
| -9 | 41 | 44 | Middle Cingulate Gyrus |
| **49** | **-5** | **-6** | **STG/Temporal Pole** |
| -7 | 25 | 42 | Middle Cingulate Gyrus |
| 43 | -37 | 14 | Inferior Frontal Gyrus |
| -15 | 71 | 10 | Calcarine Gyrus |
| 25 | 63 | -12 | Fusiform Gyrus |
| 17 | 33 | 56 | Postcentral Gyrus |
| -33 | 19 | -10 | Hippocampus |
| -29 | 63 | 18 | Middle Occipital Gyrus |
| -43 | 5 | 48 | Precentral Gyrus |
| **9** | **59** | **28** | **Precuneus** |
| 39 | -1 | 36 | Precentral Gyrus |
| 37 | 69 | -20 | Cerebellum |
| -13 | 71 | -22 | Cerebellum |
| 1 | 39 | 6 | Cerebellum |

| | | | |
|---|---|---|---|
| **33** | **19** | **-18** | **Parahippocampal Gyrus** |
| -7 | -17 | 44 | Superior Medial Gyrus |
| 13 | 77 | -22 | Cerebellum |
| 43 | 61 | 32 | Angular Gyrus |
| -63 | 17 | -6 | Middle Temporal Gyrus |
| 57 | 27 | 2 | Middle Temporal Gyrus |
| -5 | -31 | 30 | Anterior Cingulate Cortex |
| -47 | 61 | -18 | Fusiform Gyrus |
| -17 | 19 | -6 | Hippocampus |
| -53 | -7 | 34 | Precentral Gyrus |
| -9 | 83 | -2 | Calcarine Gyrus |
| -19 | 89 | 2 | Superior Occipital Gyrus |
| -37 | -43 | 12 | Middle Frontal Gyrus |
| 21 | 91 | 4 | Middle Occipital Gyrus |
| 31 | 5 | 54 | Precentral Gyrus |
| -23 | -41 | -10 | Middle Orbital Gyrus |
| 7 | 55 | 22 | Precuneus |
| -3 | -35 | 8 | Anterior Cingulate Cortex |
| 5 | 51 | 54 | Precuneus |
| 21 | 85 | 26 | Superior Occipital Gyrus |
| -13 | 15 | 46 | Middle Cingulate Gyrus |
| -19 | 63 | -8 | Lingual Gyrus |
| -1 | 41 | 28 | Posterior Cingulate Cortex |
| -19 | 31 | 48 | Paracentral Lobule |
| 23 | 5 | 64 | Superior Frontal Gyrus |
| 43 | 45 | 44 | Inferior Parietal Lobule |
| -9 | 69 | 50 | Precuneus |
| -45 | -29 | 26 | Middle Frontal Gyrus |
| **1** | **65** | **44** | **Precuneus** |
| -43 | -7 | 22 | Inferior Frontal Gyrus |
| -41 | 65 | -8 | Inferior Temporal Gyrus |
| -37 | 79 | -8 | Inferior Occipital Gyrus |
| -7 | 49 | 48 | Precuneus |
| 5 | 59 | 6 | Lingual Gyrus |
| 47 | 55 | -8 | Inferior Temporal Gyrus |
| 15 | 73 | -16 | Cerebellum |
| **17** | **71** | **18** | **SOG** |
| -51 | 27 | 38 | Supramarginal Gyrus |
| 35 | 77 | -4 | Inferior Occipital Gyrus |
| -45 | -3 | 26 | Inferior Frontal Gyrus |
| -5 | 75 | 8 | Calcarine Gyrus |
| -27 | 79 | 26 | Superior Occipital Gyrus |
| 25 | -29 | -4 | Inferior Frontal Gyrus |

| | | | |
|---|---|---|---|
| -47 | 55 | 38 | Inferior Parietal Lobule |
| **-55** | **41** | **20** | **STG** |
| -33 | -3 | 54 | Middle Frontal Gyrus |
| 35 | 47 | 40 | Inferior Parietal Lobule |
| **1** | **65** | **30** | **Precuneus** |
| -3 | 47 | -14 | Cerebellum |
| 39 | -39 | 18 | Inferior Frontal Gyrus |

18 supra-threshold regions identified via significant interaction between stimulation condition and demand (bolded), followed by their drivers—regions showing significant interaction between stimulation condition and demand in their seed-based connectivity to one of the 18 supra-threshold regions.

**Supplementary Table 4: Findings of the task-dependent connectivity analysis and their corresponding drivers for PFC-targeted stimulation.**

| Cluster Peak (RAI) | | | |
|---|---|---|---|
| x | y | z | Region |
| **-7** | **27** | **44** | **Middle Cingulate Cortex** |
| 41 | 7 | 42 | Precentral Gyrus |
| -37 | -19 | -16 | Superior Temporal Gyrus/Temporal Pole |
| 59 | -5 | 18 | Inferior Frontal Gyrus |
| -1 | -53 | 10 | Superior Medial Gyrus |
| 55 | 15 | 12 | Superior Temporal Gyrus |
| 33 | 59 | 34 | Angular Gyrus |
| -35 | 47 | -18 | Fusiform Gyrus |
| -45 | 15 | 52 | Precentral Gyrus |
| -1 | 65 | 20 | Cuneus |
| 45 | -19 | -12 | Superior Temporal Gyrus/Temporal Pole |
| -25 | 19 | 66 | Precentral Gyrus |
| -53 | 9 | 32 | Postcentral Gyrus |
| 39 | 3 | 6 | Insula |
| -15 | 81 | -6 | Lingual Gyrus |
| -63 | 33 | 4 | Middle Temporal Gyrus |
| 23 | -13 | 40 | Middle Frontal Gyrus |
| 11 | 53 | -14 | Cerebellum |
| -21 | -19 | 44 | Middle Frontal Gyrus |
| 57 | 3 | -6 | Superior Temporal Gyrus |
| 47 | 47 | 26 | Supramarginal Gyrus |
| -37 | -35 | -2 | Inferior Frontal Gyrus |
| **-5** | **61** | **-28** | **Cerebellar Vermis** |
| -15 | 35 | 40 | Middle Cingulate Gyrus |
| 1 | 67 | 6 | Lingual Gyrus |
| -15 | -21 | 44 | Superior Frontal Gyrus |
| 17 | -55 | 30 | Superior Frontal Gyrus |
| -49 | 45 | 22 | Supramarginal Gyrus |
| 23 | 41 | 62 | Postcentral Gyrus |
| -9 | -13 | 60 | SMA |
| -3 | -45 | -4 | Middle Orbital Gyrus |
| 57 | 61 | 16 | Middle Temporal Gyrus |
| -55 | 7 | -6 | Superior Temporal Gyrus |
| **25** | **-33** | **0** | **Inferior Frontal Gyrus** |
| 55 | 47 | 30 | Supramarginal Gyrus |
| **-25** | **-11** | **-2** | **Putamen** |
| -21 | 77 | -28 | Cerebellum |

| | | | |
|---|---|---|---|
| -23 | 39 | 52 | Postcentral Gyrus |
| -11 | -59 | 30 | Superior Medial Gyrus |
| 37 | -21 | 0 | Inferior Frontal Gyrus |
| -3 | -53 | 16 | Superior Medial Gyrus |
| -53 | -7 | -2 | Superior Temporal Gyrus/Temporal Pole |
| -7 | -5 | 12 | Caudate Nucleus |
| 15 | 63 | -32 | Cerebellum |
| -27 | 1 | 58 | Superior Frontal Gyrus |
| -17 | 79 | 0 | Calcarine Gyrus |
| -39 | -17 | -24 | Medial Temporal Pole |
| 35 | -19 | 46 | Middle Frontal Gyrus |
| 37 | -13 | -26 | Medial Temporal Pole |
| 13 | 1 | 18 | Caudate Nucleus |
| 49 | 25 | 28 | Supramarginal Gyrus |
| -45 | 45 | -6 | Inferior Temporal Gyrus |
| -53 | 15 | 26 | Supramarginal Gyrus |
| 49 | 1 | 38 | Precentral Gyrus |
| -39 | 63 | 30 | Angular Gyrus |
| 31 | 69 | 46 | Superior Parietal Lobule |
| 59 | 31 | -10 | Middle Temporal Gyrus |
| 39 | 55 | 28 | Angular Gyrus |
| -33 | 33 | 50 | Postcentral Gyrus |
| -29 | 17 | 56 | Superior Frontal Gyrus |
| 9 | 9 | 52 | SMA |
| 27 | 49 | -22 | Cerebellum |
| -53 | 1 | 38 | Precentral Gyrus |
| 17 | -7 | 56 | Superior Frontal Gyrus |
| -31 | 23 | 30 | Postcentral Gyrus |
| 43 | 49 | -18 | Fusiform Gyrus |
| 43 | -23 | -4 | Inferior Frontal Gyrus |
| **-13** | **-43** | **-4** | **Middle Occipital Gyrus** |
| 27 | 41 | -18 | Fusiform Gyrus |
| -47 | 55 | -8 | Inferior Temporal Gyrus |
| -21 | -1 | 10 | Putamen |
| -27 | -45 | 26 | Middle Frontal Gyrus |
| -39 | 61 | -24 | Cerebellum |
| -23 | 1 | 8 | Putamen |
| -1 | 7 | 38 | Middle Cingulate Gyrus |
| 55 | 21 | 22 | Postcentral Gyrus |
| -7 | 59 | 48 | Precuneus |
| 15 | 63 | 52 | Superior Parietal Lobule |
| **-31** | **77** | **22** | **ACC/Orbital Gyrus** |

| | | | |
|---|---|---|---|
| 51 | -5 | 14 | Inferior Frontal Gyrus |
| 51 | 63 | 8 | Middle Temporal Gyrus |
| -61 | 37 | 26 | Supramarginal Gyrus |
| -7 | 25 | 2 | Thalamus |
| 13 | -53 | 2 | Superior Orbital Gyrus |
| -5 | -25 | 38 | Middle Cingulate Gyrus |
| 3 | 43 | 50 | Precuneus |
| **-13** | **-43** | **24** | **Insula/Inferior Frontal Gyrus** |
| -29 | -15 | 46 | Middle Frontal Gyrus |
| **35** | **-21** | **4** | **ACC** |
| 49 | 7 | 28 | Precentral Gyrus |
| 29 | -27 | -10 | Inferior Frontal Gyrus |
| 7 | -25 | 14 | Anterior Cingulate Cortex |
| -49 | 51 | -22 | Inferior Temporal Gyrus |
| 35 | 9 | 0 | Insula |
| 39 | -7 | 36 | Middle Frontal Gyrus |
| 47 | 53 | -16 | Inferior Temporal Gyrus |
| 33 | 41 | 12 | Superior Temporal Gyrus |
| 5 | 31 | 42 | Middle Cingulate Gyrus |
| **-23** | **39** | **-4** | **Parahippocampal Gyrus** |
| -27 | -37 | -4 | Middle Orbital Gyrus |
| -7 | 51 | 58 | Precuneus |
| -37 | 55 | -38 | Cerebellum |
| **33** | **3** | **-22** | **Parahippocampal Gyrus** |
| -43 | -29 | 28 | Inferior Frontal Gyrus |
| -13 | -45 | -8 | Superior Orbital Gyrus |
| 45 | 65 | 32 | Angular Gyrus |
| 23 | 73 | 44 | Superior Parietal Lobule |
| -27 | -55 | 10 | Middle Frontal Gyrus |
| 25 | -21 | 50 | Middle Frontal Gyrus |
| -31 | 27 | 16 | Heschls Gyrus |
| -67 | 27 | 2 | Middle Temporal Gyrus |
| 33 | 11 | -10 | Hippocampus |
| -47 | 13 | 26 | Postcentral Gyrus |
| 15 | 49 | -20 | Cerebellum |
| 9 | 33 | 4 | Hippocampus |
| -63 | 17 | -8 | Middle Temporal Gyrus |
| -9 | 21 | 42 | Middle Cingulate Gyrus |
| -33 | -7 | 36 | Middle Frontal Gyrus |
| 7 | -25 | 32 | Superior Medial Gyrus |
| -29 | 13 | 8 | Putamen |
| -59 | -1 | 12 | Rolandic Operculum |

| | | | |
|---|---|---|---|
| 1 | 11 | 66 | SMA |
| 51 | -11 | 16 | Inferior Frontal Gyrus |
| -53 | 33 | 6 | Middle Temporal Gyrus |
| -49 | 27 | 22 | Rolandic Operculum |
| -51 | 15 | 28 | Supramarginal Gyrus |
| -33 | -35 | 20 | Middle Frontal Gyrus |
| -37 | 59 | 32 | Angular Gyrus |
| **-29** | **59** | **32** | **ACC** |
| 39 | -23 | -6 | Inferior Frontal Gyrus |
| 57 | 41 | 26 | Supramarginal Gyrus |
| 7 | 69 | 46 | Precuneus |
| -1 | -43 | 40 | Superior Medial Gyrus |
| **-35** | **-39** | **2** | **Inferior Frontal Gyrus** |
| 59 | 5 | -2 | Superior Temporal Gyrus |
| 1 | -31 | 54 | Superior Medial Gyrus |
| -47 | 43 | -2 | Middle Temporal Gyrus |
| 45 | 21 | 24 | Supramarginal Gyrus |
| 47 | 61 | -22 | Cerebellum |
| **-35** | **55** | **16** | **Middle Occipital Gyrus** |
| -47 | 65 | -18 | Cerebellum |
| -33 | 85 | -4 | Inferior Occipital Gyrus |
| 27 | 63 | 52 | Superior Parietal Lobule |
| -25 | 65 | 50 | Superior Parietal Lobule |
| -21 | 61 | 10 | Calcarine Gyrus |
| -47 | 9 | 44 | Precentral Gyrus |
| **-7** | **-23** | **28** | **Orbital Gyrus** |
| 41 | 7 | 40 | Precentral Gyrus |
| -37 | 47 | -22 | Cerebellum |
| -25 | -1 | 8 | Putamen |
| **-15** | **-7** | **-10** | **Superior Temporal Gyrus** |
| 49 | -5 | 14 | Inferior Frontal Gyrus |
| 41 | -29 | -4 | Inferior Frontal Gyrus |

15 supra-threshold regions identified via significant interaction between stimulation condition and demand (bolded), followed by their drivers—regions showing significant interaction between stimulation condition and demand in their seed-based connectivity to one of the 15 supra-threshold regions.
